# Mitonuclear effects on sex ratio persist across generations in interpopulation hybrids

**DOI:** 10.1101/2023.09.08.556888

**Authors:** Suzanne Edmands, Jacob R. Denova, Ben A. Flanagan, Murad Jah, Scott L. Applebaum

## Abstract

Eukaryotic energy production requires tight coordination between gene products from both the nuclear and mitochondrial genomes. Because males and females often have different energetic strategies, this mitonuclear coordination can be expected to differentially impact the two sexes. Previous work found evidence for sex-specific mitonuclear effects in the copepod *Tigriopus californicus* by comparing two parental lines and their reciprocal F1 crosses. However, an alternative hypothesis is that the patterns could instead be driven by the parental source of nuclear alleles. Here we test this alternative hypothesis by extending the same cross to F2 hybrids, who receive both maternal and paternal nuclear alleles from F1 hybrids. Results confirm mitonuclear effects on sex ratio, with distorted ratios persisting from the F1 to F2 generations, despite reduced fitness in F2 hybrids. No sex by cross interactions were found for other phenotypic traits measured. Mitochondrial DNA content was shown to be higher in females, the more stress-tolerant sex. Both routine metabolic rate and oxidative DNA damage were found to be lower in F2 hybrids than in parentals. Confirmation of sex-biased mitonuclear effects in *T. californicus* is notable, given that the species lacks sex chromosomes, which can confound interpretations of sex-specific mitochondrial effects.

## 1. Introduction

In the transition from free-living microbe to captured organelle, mitochondria transferred much of their genome to the nucleus, leaving the typical mitochondrial genome with only 13 protein-coding genes, 2 rRNA genes and 22 tRNA genes^1^. Mitochondrial function thus relies on a very large number of nuclear genes (∼1500) whose products function in the mitochondria^2,3^. Some of these nuclear-encoded genes form tight complexes with mitochondrial-encoded genes and these chimeric complexes are crucial to the production of energy through oxidative phosphorylation (OXPHOS), as well as to the replication, transcription and translation of the mitochondrial genome^3,4^.

Because interactions between the products of mitochondrial DNA (mtDNA) and nuclear DNA (nDNA) are central to metabolism, these mitonuclear interactions might be expected to have different effects in males and females, who often have quite different metabolic demands^5,6,7^. Indeed, even if the two sexes have equivalent metabolic rates, they may have dramatically different energy allocations to different tissue types or life history components^8,9,10^. Selection on mtDNA may also be sex-biased due to the asymmetrical inheritance of mtDNA. In the majority of bilaterians, mitochondria are inherited maternally, making males a dead-end for mitochondrial transmission. The absence of direct selection on male mitotypes has been proposed to result in a “mother’s curse” in which males accumulate deleterious mtDNA load^11^. A subset of studies, including particularly rigorous studies in *Drosophila*, find strong support for this phenomenon^e.g. 12,13,14,15^. Other studies find sex-specific mitochondrial and mitonuclear effects, and yet do not find them to be consistently more deleterious in males^e.g. 16,17,18^.

Our understanding of the ubiquity of sex-specific mitonuclear effects is impeded by the paucity of studies reporting results for both sexes. This is a well-recognized issue in the biomedical literature, where women were historically excluded from studies to reduce variation from hormonal cycles and to protect female reproduction^19,20^. Beyond the medical literature, ecological studies also frequently fail to address sex differences. For example, less than half of stress tolerance studies were found to report results for both sexes^21^. Reasons for these omissions include one sex being more abundant or easier to collect, difficulties in distinguishing sexes, and a focus on early-life stages.

Poor understanding of how mitonuclear effects differ between sexes is problematic because they have important medical implications. Mitochondrial dysfunction has long been associated with age-related decline^22,23^ and aging phenotypes are often sex-specific^24,25,26^. Understanding the extent of sex-specific mitochondrial effects will be critical for assessing the need for sex-specific therapies. This is particularly true for mitochondrial replacement therapy, a potential remedy for heritable mitochondrial diseases. It has been suggested that potential risks of this therapy could be limited by restricting it to male embryos who will not pass donor mtDNA onto the next generation^27,28^, yet this strategy may be ill-advised if mtDNA effects are indeed more detrimental to males. Beyond medical implications, sex-specific mitonuclear effects also have broad implications for the maintenance of genetic variation. Modelling work shows that sex differences in mitonuclear effects promote stable mtDNA polymorphisms within populations^29,30,31,32^.

In this study we tested for sex-specific mitonuclear effects in the marine copepod *Tigriopus californicus*. This species is becoming a model for understanding mitonuclear coevolution^33,34,35,36,37,38,39^ with the distinct advantage that viable and fertile hybrids can be created between populations with highly divergent mitotypes^40,41^. *T. californicus* does not have sex chromosomes (instead, sex determination is polygenic^42,43,44,45^), which means that sex-specific mitochondrial effects are not confounded with sex chromosome effects. Despite the absence of sex chromosomes in this species, males and females show substantial differences in stress tolerance^46,47,48,49,50^ and gene expression^51,52^.

In previous work in this species^53^ we tested for sex-specific mitonuclear effects in a cross between highly genetically divergent populations from California and Washington, USA. These two populations are ∼3% divergent across the nuclear genome and ∼20% divergent for mtDNA, with differentiation across all 37 mitochondrial genes^37,18^. Results showed differences in sex ratio and sex-specific survivorship between reciprocal F1 hybrids^53^. Since these F1 cohorts have different mitochondrial haplotypes on the same 50:50 nuclear background, this finding is consistent with sex-specific mitonuclear interactions. However, an alternative explanation is that these differences are due to the parental source of nuclear alleles with, for example, one F1 receiving Californian nuclear alleles from its mother, and the reciprocal F1 receiving Californian nuclear alleles from its father. This could lead to sex differences between the F1 cohorts if, for example, there is genomic imprinting wherein genes inherited from male vs. female parents are differentially expressed due to epigenetic modifications^54^.

Here we test this alternative hypothesis using the same pair of populations and extending hybridization to the F2 generation. If sex differences between reciprocal F1 are due to the parental source of nuclear alleles, these differences should be erased in the F2, who receive both maternal and paternal nuclear alleles from F1 hybrids. As in the prior study^53^ we assayed parental and hybrid lines for offspring numbers, adult sex ratio, and sex-specific mtDNA content and DNA damage. In addition, lines were assayed for routine metabolic rate. Extending this analysis to second generation hybrids also allows us to test how sex-specific phenotypes are impacted by increased mitonuclear mismatch.

## 2. Materials and methods

### (a) Population sampling and culture maintenance

Animals were collected from supralittoral pools at Friday Harbor Laboratories, WA, USA (F, 48.55N 117.25) and San Diego, CA, USA (S, 32.74N 117.25W; figure 1). All animals were maintained in incubators set at 20°C with a 12h light: 12h dark cycle and fed ground Spirulina (Nutraceutical Science Institute, USA) and ground Tetramin fish food (Tetra Holding Inc., USA). Periodically, cultures were rehydrated with diH_2_O and partial seawater changes were conducted. Seawater in this study was obtained from the USC Wrigley Marine Science Center (Santa Catalina Island, CA, USA) and filtered 3x with a 37μm filter. Inbred lines were created by full-sibling mating for a minimum of 50 generations before experimental crosses began. Notably, these are the same inbred lines used in the previous study^53^ of sex-specific mitonuclear effects, but with approximately 40 additional generations of inbreeding.

**Figure 1.**
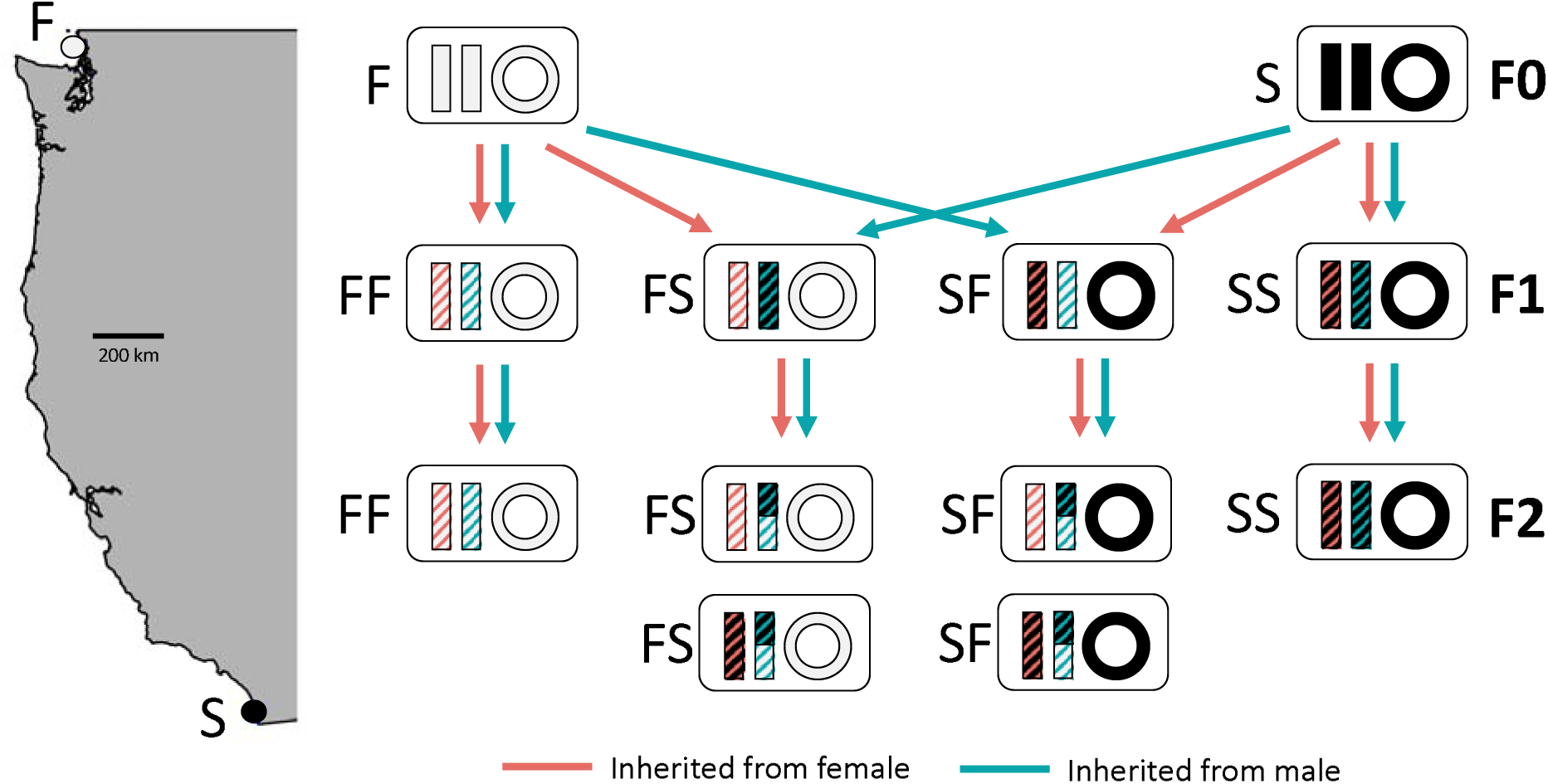
Experimental cross design. Inbred lines from population F (Friday Harbor Laboratories, WA) and S (San Diego, CA) were crossed to produce F1 cohorts FF, FS, SF and SS. Each of these F1 cohorts was then crossed to itself to produce the F2 cohorts. Bars indicate nuclear chromosomes and open circles indicate mitochondrial genomes. Because meiosis is achiasmatic in females of this species^55,56^, only males pass recombinant chromosomes. All cross names are shown as female x male.

### (b) Experimental crosses, offspring numbers and sex ratio

Inbred lines F and S were first sequenced for a segment of mitochondrial 12s (forward primer 5’-CCAAGATACTTTAGGGATAACAGC-3’, reverse primer 5’-CTGTTCTATCAAAACAGCCCT-3’) to confirm there had been no contamination. Crosses (figure 1) were established between lines F and S to produce F1 cohorts FF, FS, SF and SS (all crosses listed as female x male). Notably, F1 hybrids FS and SF have mitochondria from alternative parents and nuclear genomes that are alike in having a 50:50 F:S ratio, but different in that nuclear alleles are inherited from different parents. Each of the four F1 cohorts was then crossed with itself to produce F2 cohorts FF, FS, SF and SS. Because each F1 cohort acts as both mother and father, the FS and SF F2 cohorts thus have equivalent nuclear genotypes but different mitochondrial haplotypes.

All experimental crosses were conducted in Petri dishes with culture medium containing 0.1g Spirulina and 0.1g Tetramin per liter of triple filtered seawater. In this species, mature males clasp virgin females and guard them until the female reaches reproductive maturity. The male then inseminates the female who mates only once and uses stored sperm to fertilize multiple clutches of eggs. To obtain males and virgin females we placed each clasped pair onto damp filter paper under a dissecting microscope and separated the two animals using a fine probe. F0 crosses for each cohort (FF, FS, SF and SS) were generated from sets of 5 females and 5 males in each dish. Once a week, dishes were rehydrated, fed concentrated food (0.04g Spirulina and 0.04g Tetramin in 4ml seawater) and observed under a dissecting microscope. When F0 females formed egg sacs they were moved to their own new dish. When larvae were observed, mothers were again moved to a separate dish. F0 mothers were retired after producing a maximum of three sets of larvae (that is, a maximum of three larval cohorts per family). F1 offspring dishes were allowed to mature for 28d after offspring were first observed. On day 28, male and female F1 offspring were enumerated and up to two gravid females were each moved to their own new dish to produce F2 offspring.

Once a week, these F1 female dishes were rehydrated, fed and monitored in the same way as the F0 female dishes, and F1 mothers were retired after producing a maximum of three sets of offspring. F2 offspring dishes were allowed to mature for 28d after larvae were first observed. On day 28, females were counted and moved into one new dish, and males were counted and moved into a different dish. On day 56 after larvae were first observed, surviving females and males were re-counted. F2 animals were then either used for metabolic rate assays or frozen at -80°C for subsequent assays of mtDNA content and oxidative DNA damage. F1 animals were not used for these additional assays because they were needed as parents for the F2 generation.

Data were analyzed using R v. 4.1.^57^. Cases where dams produced zero offspring were omitted from analysis. Offspring numbers and survivorship were treated as non-parametric. Effects of cross and sex were assessed by Kruskal-Wallis tests, with pairwise comparisons calculated by Wilcoxon’s tests with adjusted *p*-values.

### (c) mtDNA content estimation

Mitochondrial copy number was estimated in animals frozen at day 56 after larvae were first observed, using methods based on Flanagan et al^53^. DNA was extracted from individual copepods by incubation for 1 h at 65°C in 50 μl proteinase K (200 μg ml^-1^) cell lysis buffer (10mM TRIS, 50mM KCl, 0.5% Tween 20, at pH 8.8) followed by 15 min denaturation at 100°C. MtDNA copy number was estimated through quantitative polymerase chain reaction (qPCR) targeting the single copy genes *AtpC* (nuclear) and *Atp6* (mitochondrial) using previously-designed^53^ primers (*AtpC*-F 5’-CCAAGTTCATCGGAGCTGGT-3’; *AtpC*-R 5’-TACGGGCGTAACCGATGATG-3’; *Atp6*-F 5’-TGAGAACCAGAATGAACGGCT-3’; *Atp6*-R 5’-AGGGTCTTCTCGTCCCTGAA-3’). Primer annealing temperatures were optimized by gradient PCR, and we followed amplification with a melt curve and agarose gel to ensure each primer pair generated a single amplicon. To generate the standard curves for each primer pair, we performed four, five-fold serial dilutions on DNA extracted from pooled animals. The efficiencies of each primer set were 90–110% with r^2^ > 0.95. The qPCR reaction mixture consisted of 1X Brilliant III Ultra-Fast SYBR® Green QPCR Master Mix (Agilent), 0.002X reference dye (Agilent), 0.5 μM Primers and 2 µl DNA lysate. Reaction conditions were as follows: 95°C for 3 min for initial denaturation step, followed by denaturation at 95°C for 15 s, annealing at 65 for the nuclear target and 55 for the mitochondrial target for 15 s, and extension at 60°C for 15 s for the nuclear target and 30 s for the mitochondrial, repeated 40 times. qPCR reactions were run on an Agilent AriaMx Real-Time PCR System (Agilent), and threshold values (Ct) were obtained using the accompanying software regression. Each reaction was performed in triplicate and the mean was used to calculate mtDNA content in a delta-Ct manner according to Rooney et al^58^. In cases where the triplicate values were not within 1 Ct value they were reassayed until the three replicates fell within 1 Ct.

Results were analyzed using R v. 4.1.3^57^. Data were log transformed to meet normality assumptions. Effects of cross, sex and their interactions were assessed by ANOVA.

### (d) Metabolic rate assays

Routine metabolic rate was estimated for individual adult F2 females and males 60-70d after larvae were first observed. Animals were transferred to autoclaved seawater without food the night before assays began. Oxygen consumption rates were measured with an optical microplate respirometer (24-well Microplate Core system, 80 μl wells, Loligo Systems, Viborg, Denmark).

To begin assays, individual copepods (both males and females) were briefly placed on filter paper and egg sacs were removed from females using a fine probe. Each individual was then transferred to a well containing autoclaved and 0.2 μm filtered air-saturated seawater. For each 24-well plate, two wells contained only filtered seawater (no copepod) to control for any non-copepod oxygen consumption. After sealing wells, oxygen consumption was recorded at 5 min intervals and an initial 25 min period was allowed for equilibration of sealed wells. Oxygen consumption rates were determined as the slope of the decline in oxygen concentration over 185 min, beginning after the initial 25 min equilibration phase. Assays were done at 23°C with the respirometer in darkness to eliminate impacts of ambient light on optical measurements. Oxygen concentration remained in the normoxic range throughout the assays. When assays were completed, wells were opened, and copepods inspected for mortality. Copepods were then collected for subsequent measurements of body length and total protein content to account for the influence of variation in size on metabolic rates.

For body length measurements, each copepod was transferred to a microscope slide and oriented for a dorsal, size-calibrated photomicrograph. Images were taken with a digital camera (Canon EOS Rebel SL3, Canon USA) attached to a stereomicroscope (Leica M-Series). The total length of each copepod (from anterior of eye spot to fork in caudal furca) was measured using an ocular micrometer and ImageJ software^59^. Total protein content of individual copepods was determined using a bicinchronic acid (BCA) microassay (Pierce Micro BCA Protein Assay, Thermo Fisher Scientific). Tissues were disrupted by adding 350 μl ultrapure water and sonicating for 10s at 7 watts RMS. The sample was held on ice while sonicating to avoid heating and denaturing proteins. Two replicate subsamples of the homogenate (150 μl each) were transferred to clean test tubes and assayed according to manufacturer’s instructions, with a total assay volume of 300 μl. Samples were then incubated for 60 min at 60°C. A standard curve was created using the bovine serum albumin standard provided by the manufacturer. After incubation, absorbance for each sample and protein standard was measured at 562 nm in a SpectraMax plate reading spectrophotometer (Molecular Devices, LLC, San Jose, CA). The standard curve was used to convert absorbances to total protein content (μg ind^-1^).

Data were analyzed using R v. 4.1.3^57^. There was no mortality during the respiration assays, but two out of 88 individuals assayed had no detectable respiration above the no animal control measurements. These were removed from the metabolic rate dataset. To meet normality assumptions for statistical tests, data for body length, protein content, and protein-specific respiration were log transformed. Effects of cross, sex and their interactions were assessed by ANOVA.

### (e) Oxidative DNA damage assays

DNA damage caused by oxidative stress was estimated by measuring 8-hydroxy-2′-deoxyguanosine (8-OH-dG) content, which specifically quantifies DNA damage resulting from guanosine oxidation. Samples frozen 56 d after larvae were first observed were combined by sex within each family and age class to estimate 8-OH-dG damage using enzyme-linked immunosorbent assay (ELISA; Cayman Chemical cat. 589320). The minimum amount of DNA required for the ELISA reaction was determined to be 5 ng. To maximize DNA extraction yield, we used a phenol-chloroform extraction technique. First, individuals underwent proteinase K-lysis buffer extraction (see *mtDNA content estimation*). Then 100 μL of the phenol-chloroform mixture (pH 8.0: VWR) was added to each lysate and the mixture was vortexed and then centrifuged for 5 min. The aqueous layer was removed and an additional 80 μL of 0.5x TE (pH 8.0) was added and samples were briefly vortexed and then centrifuged for 5 min. The second aqueous layer was combined with the first and 500 μL ice-cold 95% ethanol, 1 μL GlycoBlue (Invitrogen), and 78 μL of 3 M NaOAc. The samples were inverted and incubated at -20°C for 1 h and then centrifuged for 30 min. The ethanol was decanted leaving the pellet which was washed again with 300 μL ice-cold 70% ethanol and centrifuged for 10 min. The ethanol was again decanted, and the remaining DNA was dried by vacuum centrifuge. DNA samples were resuspended in 60 μL molecular grade water and DNA was quantified using Qubit™ 3 Fluorometer (Invitrogen) using Qubit™ dsDNA HS Assay Kit (Invitrogen). Samples were then treated with P1 nuclease (New England Biolabs) then rSAP (New England Biolabs), and three replicate individuals were pooled by sex within each family and age. 8-OH-dG damage ELISA was performed according to the manufacturer’s protocol. For each sample, four technical replicates were split between two concentrations of DNA and averaged, and data were analyzed as the ratio of DNA damage to total DNA.

ANOVA was performed in R v. 4.1.3^57^ to estimate the effects of cross, sex and their interactions on DNA damage. *Post-hoc* testing was done using least-square means with Tukey adjusted *p*-values.

## 3. Results

### (a) Offspring numbers and sex ratio

The total number of F1 offspring reaching adulthood (figure 2a) showed a significant effect of cross (Kruskal Wallis, *p* = 0.005) with a moderate effect size (Eta squared, η^2^) of 0.066. While the mean number of offspring in both hybrid cohorts was above parentals, pairwise comparisons (Wilcoxon’s) showed significant differences only between FF and FS (*p* = 0.002, effect size (Z statistic) = 0.422). In the F2 generation (figure 2b), cross was again significant (Kruskal Wallis, *p* = 1.34e^-9^) with a large effect size (Eta squared, η^2^) of 0.148. Pairwise comparisons (Wilcoxon’s) showed both hybrid cohorts significantly below both parental cohorts, with the smallest difference between FF and FS (*p* = 0.031, effect size (Z statistic) = 0.242) and the largest between SF and SS (*p* = 6.42e^-9^, effect size (Z statistic) = 0.466). The proportion of F2 offspring surviving from d28 to d56 (electronic supplementary material, figure S1) showed a significant effect of cross (Kruskal Wallis, *p* = 0.004) with a small effect size (Eta squared, η^2^) of 0.016. Pairwise Wilcoxon tests found survival in the FF parental was lower than the FS hybrid (*p* = 0.044, effect size (Z statistic) 0.215) and the SS parental. (*p* = 0.002, effect size (Z statistic) 0.260). Effects of sex were not significant (Kruskal Wallis, *p* = 0.415).

**Figure 2.**
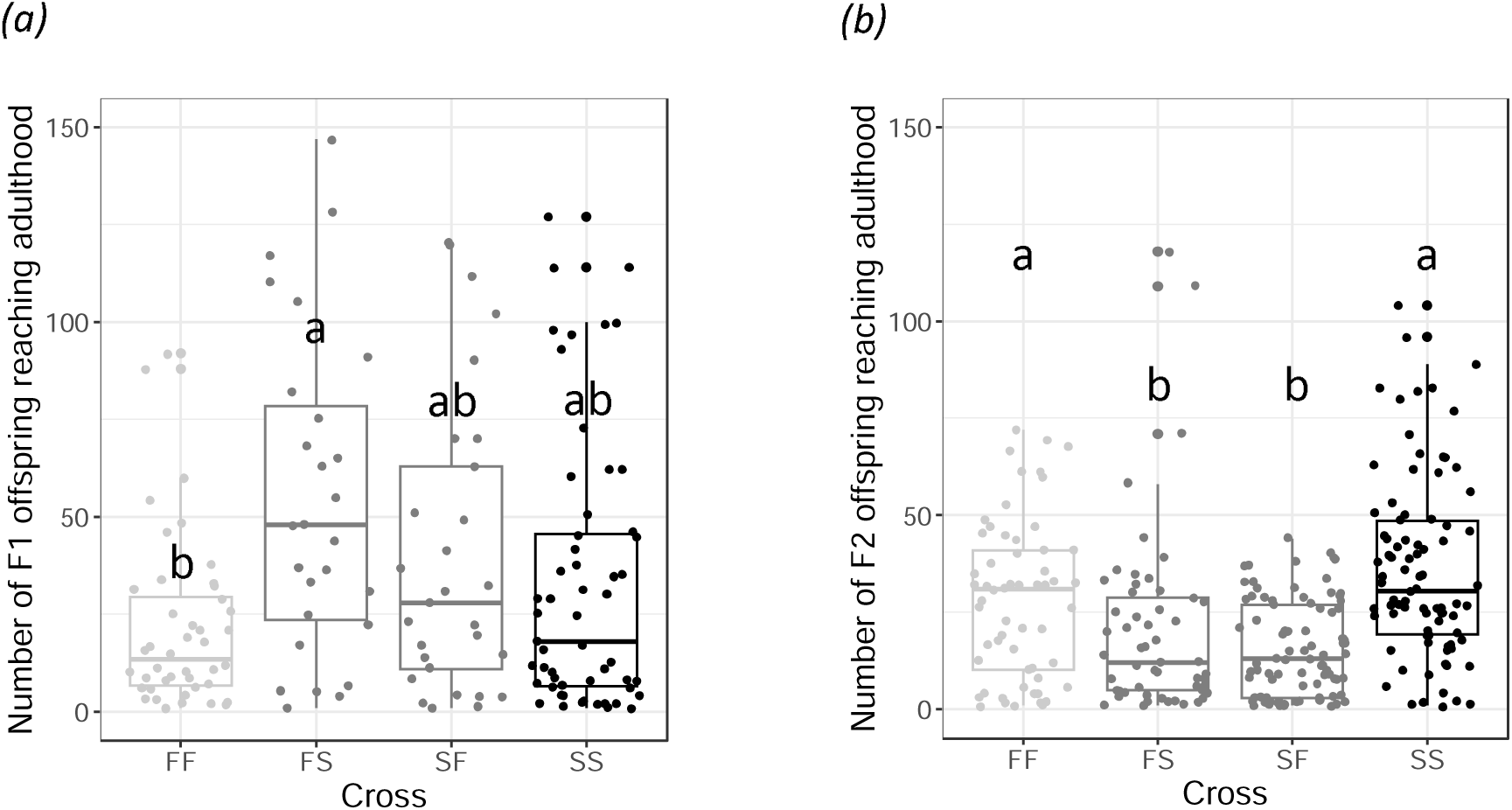
Number of offspring surviving to adulthood (per family per week) in F1 (a) and F2 (b). Letters above bars denote differences between cross types from pairwise Wilcoxon tests. *N* = 49-77 observations from 23-31 families for each cross in the F1 (a) and 62-91 observations from 25-41 families for each cross in the F2 (b).

Sex ratio in the F1 (figure 3a), measured as the number of each sex at day 28, showed a significant excess of males in FS hybrids (Wilcoxon test, *p* = 0.0094, effect size (Z statistic) = 0.372). In the F2 at day 28 (figure 3b), a significant excess of males was found in both FS hybrids (*p* = 0.029, effect size = 0.216) and SS parentals (*p* = 6.3e^-6^, effect size = 0.349). When F2 offspring were recounted at day 56 (electronic supplementary material, figure S2) these same patterns persisted, with excess males in FS (*p* = 0.0077, effect size = 0.312) and SS (*p* = 0.00095, effect size = 0.307).

**Figure 3.**
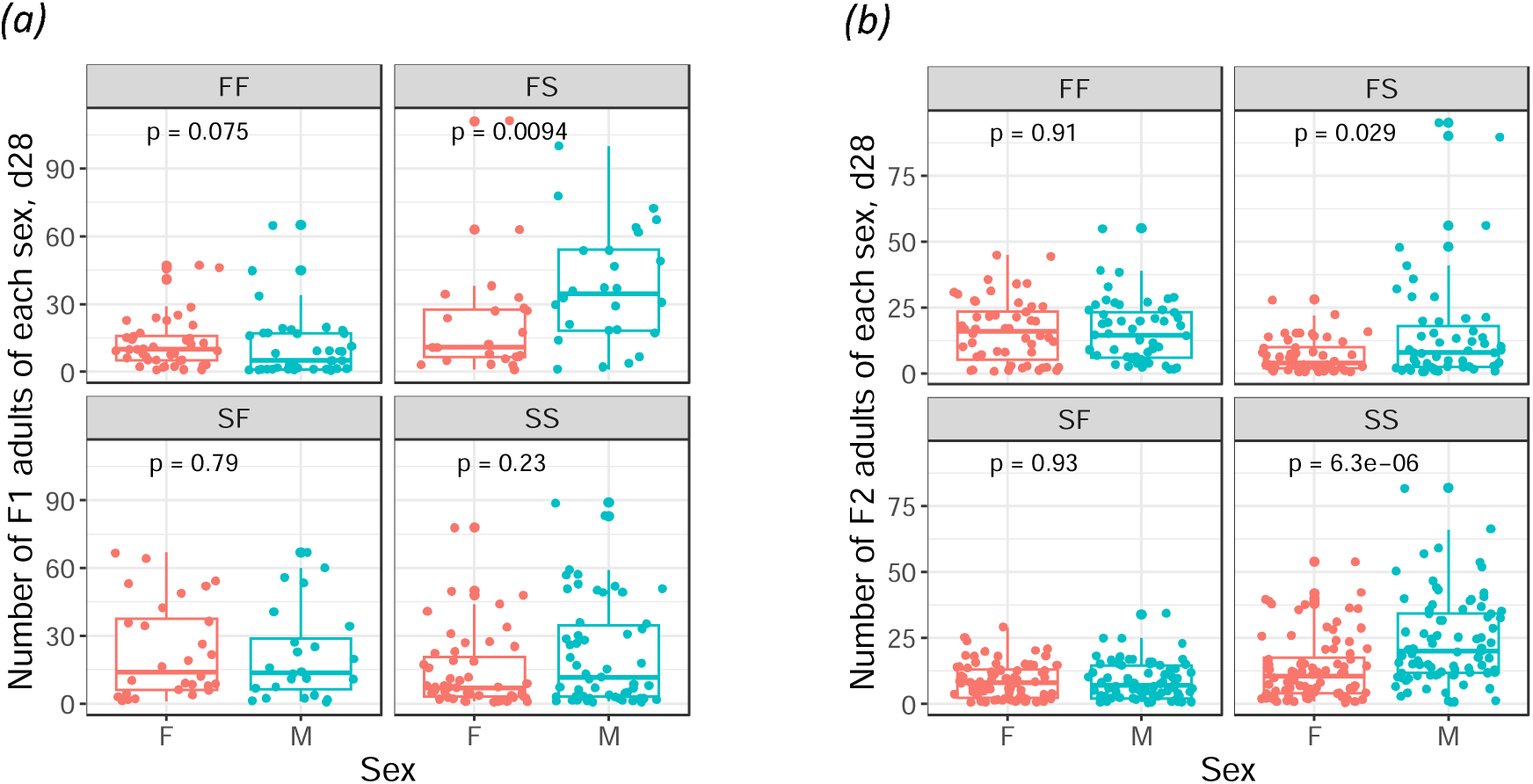
Sex ratio, measured as the number of females (F) and males (M) surviving to adulthood (per family per week) in F1 (a) and F2 (b). *P*-values indicate sex differences by pairwise Wilcoxon tests. *N* = 49-77 observations from 23-31 families for each cross in the F1 (a) and 62-91 observations from 25-41 families for each cross in the F2 (b).

### (b) mtDNA content

Log transformed mtDNA content data was used for analysis of variance on sex, cross, and their interactions. Only sex was significant (*p* = 0.00186, with a moderate effect size of 0.22 by partial Eta squared (η_p_^2^)), with mtDNA content higher in females than in males (figure 4).

**Figure 4.**
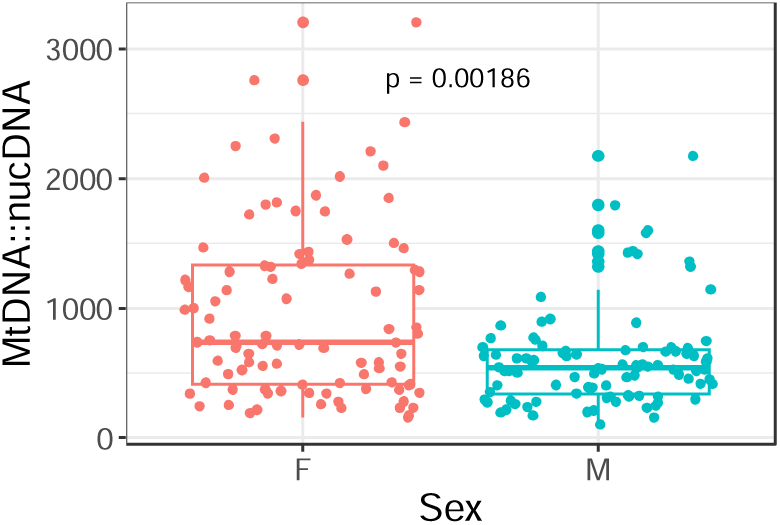
Mitochondrial DNA content for F2 females (F) and males (M) at day 56, with crosses combined. *P*-value indicates effect of sex in a two-way ANOVA on cross and sex in log transformed data. *N* = 96 individuals per sex.

### (c) Metabolic rates

Metabolic rates scale with biomass^60^ and size differences among organisms should be accounted for when comparing metabolic rates. Animals used for metabolic rate assays were each measured for body length and total protein content. Analysis of variance on log transformed body length data showed a significant interaction between cross and sex (*p* = 0.0119), with comparisons within crosses showing males being longer in the FF cross (*p* = 0.0056) but equivalent to females in the remaining 3 crosses (pairwise *t*-tests; electronic supplementary material, figure S3). Analysis of variance on log-transformed protein data showed a significant effect of sex (females 24.6% higher than males, *p* = 3.72e^-5^) as well as cross (*p* = 1.59e^-7^), with FF lower than both hybrids, SS lower than both hybrids, and SF lower than FS (Tukey HSD). Length is a convenient and easily measured value that has been used to correct for the effect of variation in size on whole animal respiration rates. This approach, however, presumes a relationship with biomass. In this study, body length and total protein content of *T. californicus* were uncorrelated (*R* = 0.042, *p* = 0.7, method = Pearson; electronic supplementary material, figure S4) indicating that length-specific metabolic rates do not effectively account for size variation. We therefore normalized metabolic rates to total protein, which constitutes the largest fraction of biochemical content in *T. californicus*^61^ and is proportional to respiring biomass.

Analysis of variance on log-transformed, protein-specific metabolic rates (metabolic rate ∼ cross * sex) revealed a significant effect of cross (*p* = 0.026) with a moderate effect size of 0.12 by partial Eta squared (η_p_^2^). Cross comparisons (figure 5) show that hybrid means are below parental means, with the SF hybrid being significantly below both parentals by Tukey HSD.

**Figure 5.**
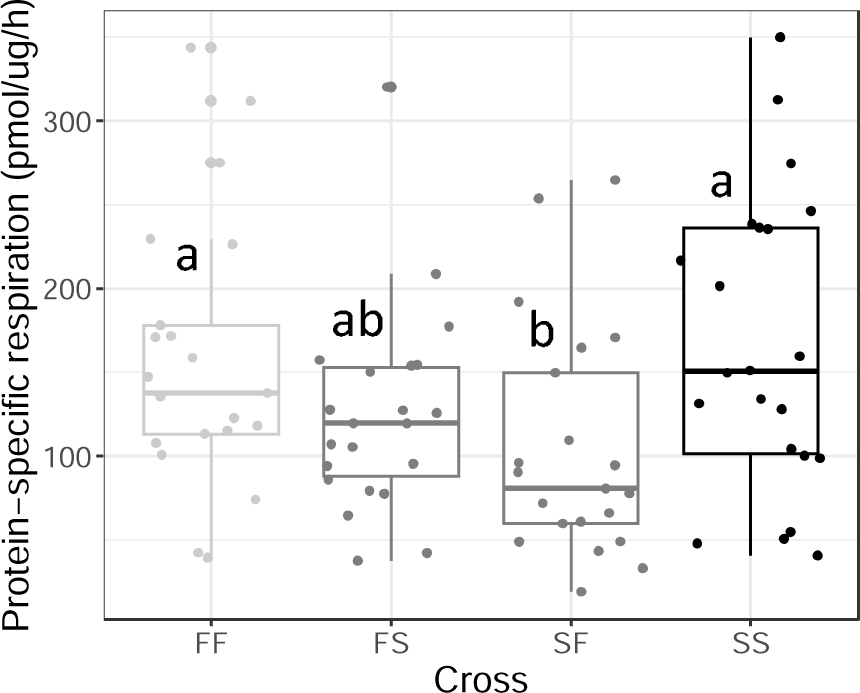
Protein-specific routine metabolic rates in F2 crosses at age 60-70 days. Letters above bars indicate differences between crosses for log transformed data in Tukey HSD tests. *N* = 21-22 individuals for each of the four crosses.

### (d) Oxidative DNA damage

DNA damage data was estimated by measuring 8-OH-dG content (pg) relative to total DNA content (μg). ANOVA on total DNA content (DNA ∼ cross * sex) found a significant effect of cross (*p* = 7.08e-5) and sex (*p* = 7.77e-9), with average DNA content being 64% higher in females. ANOVA on 8-OH-dG DNA damage (damage ∼ cross * sex) found that only cross was significant (*p* = 1.83e-5) with a large effect size of 0.46 by partial Eta squared (η_p_^2^). *Post-hoc* testing by Tukey HSD showed damage in SF hybrids lower than both parentals, and damage in FS hybrids lower than FF parentals (figure 6).

**Figure 6.**
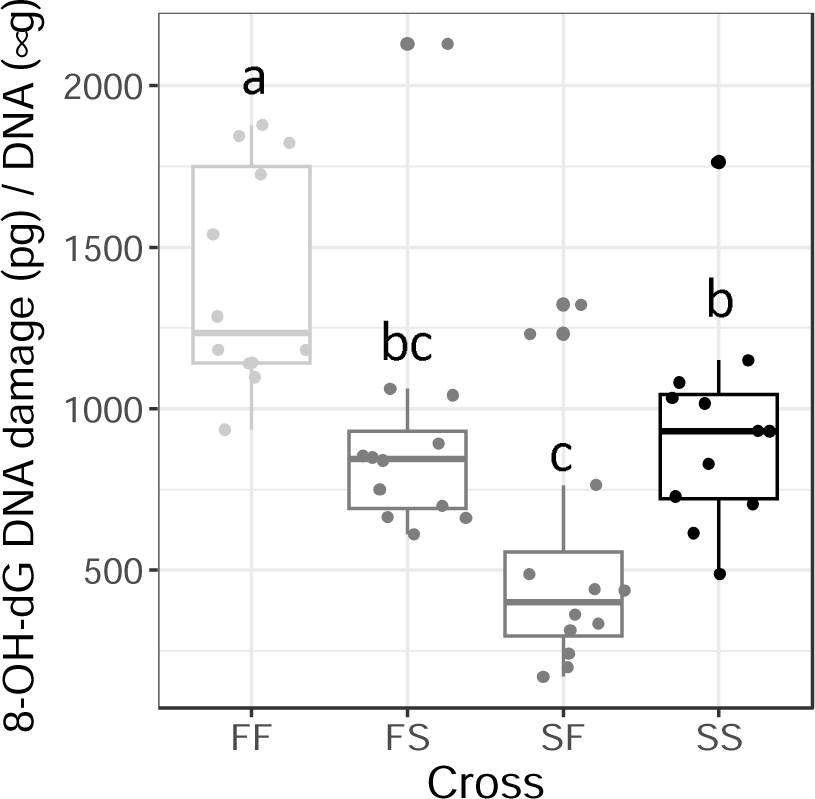
DNA damage in each F2 cross at day 56. Letters above bars indicate differences between crosses in Tukey HSD tests. *N* = 12 assays for each of the four crosses.

## 4. Discussion

Looking at both sexes combined, the total number of offspring surviving to adulthood supports the pattern of F1 heterosis and F2 hybrid breakdown commonly seen in *T. californicus* interpopulation crosses^62,41^. In the current study, both F1 cohorts were above both parentals, although not significantly so, while both F2 cohorts were significantly below both parentals. F1 heterosis is often attributed to beneficial dominance while F2 breakdown is often attributed to detrimental epistasis, including both mismatched homozygote x homozygote nuclear combinations and mismatched hemizygote x homozygote mitonuclear combinations.

Our sex ratio data for the F1 and F2 show mitonuclear effects, confirming previous results by Flanagan et al.^53^ for F1 hybrids in the same pair of populations. Like the previous study, our F1 results showed male bias in FS F1 and equivalent sex ratios in SF F1. If the difference between reciprocal F1 cohorts was due to the parental source of nuclear alleles, the difference should be erased in reciprocal F2 cohorts, who receive both maternal and paternal nuclear alleles from F1 hybrids. Instead, we found that patterns of sex bias persisted into the second generation, despite a reduction in fitness in F2 hybrids, with FS hybrids again being male biased and SF hybrids again having equivalent sex ratios. Assuming the mtDNA does not carry meiotic drive elements causing unequal allelic transmission^63^, this pattern implies that the F mitotype on a hybrid background favors male survival. Consistent with this, the sex difference between reciprocal F1 at day 28 became even stronger in animals that survived to day 56. The similar patterns of sex bias across generations suggests that sex-specific hemizygote x heterozygote mitonuclear interactions in the F1 were not overridden in the F2 by additional hemizygote x homozygote mitonuclear interactions or homozygote x homozygote nuclear-nuclear interactions. The persistence of mitonuclear effects on sex ratio across generations is important, given evidence that such interactions maintain mitochondrial polymorphism underlying trait variation^29,30,31,32^.

Mitochondrial DNA content did not show cross effects, but did show a sex effect, being higher in females than males. This is the same pattern previously found in *T. californicus*^53^ as well as in other taxa ranging from Drosophila^64^ to humans^65^. MtDNA content has been proposed as a biomarker for physiological robustness^66,67,68^ as it is positively associated with metabolic efficiency and stress tolerance. Our finding of higher mtDNA content in females is therefore consistent with prior studies in this species showing that females have higher tolerance to a range of stressors including high temperature^46,47,48^, high and low salinity^69,48^, hypoxia^50^, copper^48^, bisphenol A^48^, and paraquat^49^.

Routine metabolic rates showed a significant cross effect, but no sex effect. Mean values for F2 hybrids were below parentals, with one hybrid being significantly below both parentals. Routine metabolic rate reflects metabolism averaged over a set time interval, for animals exhibiting spontaneous behaviors (absent environmental perturbations such as stress, feeding, etc.). For ectotherms that cannot be directed or constrained to conditions for measurement of standard metabolic, RMR offers a standardized approximation of maintenance metabolism, the minimum energetic cost of sustaining cellular structure and function. Higher maintenance metabolism may be advantageous when food resources are abundant, but disadvantageous when they are scarce, since energy used for maintenance comes at the expense of energy used for growth or reproduction^70,71^. Previous work on *T. californicus* found reduced mitochondrial ATP production in hybrids^72^, which might be expected to cause a compensatory increase in oxygen consumption^73^. Indeed, studies in other taxa have found elevated metabolic rates in hybrids attributed to mitonuclear mismatch^73,74,75,76^. Our study instead found lower routine metabolic rate in hybrids, a pattern which is more rarely reported^e.g.77^. Whether this reduced maintenance metabolism increases or decreases hybrid fitness can be expected to depend on the abundance of resources.

8-OH-dG DNA damage also showed no sex effect but did show a significant cross effect. Both F2 hybrids were below both parentals, one significantly so. This is in direct contrast to previous work in the species by Barreto and Burton^78^ who found elevated oxidative DNA damage in hybrids, particularly those with the lowest fitness and greatest interpopulation divergence. Such a result is consistent with the hypothesis that mitonuclear mismatch results in a slowing of the flow of electrons in oxidative phosphorylation and an increase in free radical leak, promoting oxidative damage^79,80,81^. The contrasting results of these two studies suggest that the relationship between mitonuclear mismatch and oxidative damage is complex, and that other factors may be involved even within a single species. One factor that may contribute to the difference between studies is that the prior study used more advanced generation hybrids (F9 recombinant inbred lines, rather than F2 hybrids), with the potential for greater mitonuclear conflict. Our assays were also done on older animals (56 days, rather than 21-24 days) allowing greater time for viability selection to promote the most beneficial recombinant genotypes in the F2 hybrids. Notably, levels of DNA damage roughly parallel estimates of routine metabolic rate in our four cohorts, with SF being the lowest for both metrics, followed by FS. These lower respiration rates may contribute to the reduced DNA damage we found in hybrids.

In sum, extending a prior study^53^ to second generation hybrids revealed a number of important results. Mitochondrial DNA content was again found to be elevated in females, the more stress tolerant sex. Fitness declined in F2 hybrids, as expected, and these F2 hybrids had unexpectedly low routine metabolic rates and DNA damage. Finally, mitonuclear effects on sex ratio were confirmed, and these effects persisted from the F1 to the F2 generation, despite changes in hybrid fitness. This makes *T. californicus* one of very few species in which sex*mtDNA*nucDNA interactions have been explicitly demonstrated^e.g.82,32,31^, and perhaps the only one where they have been found in the absence of sex chromosomes which can have confounding sex-specific effects. Better understanding of the ubiquity of such 3-way interactions is needed, given the central role of mitochondria in metabolism, and the myriad possibilities for sex differences in disease and aging.

## Data accessibility

The data and code supporting this paper are available from Zenodo: ****URL and citation*****

## Authors’ contributions

S.E.: conceptualization, data curation, formal analysis, funding acquisition, investigation, project administration, supervision, validation, visualization, writing-original draft, writing-review & editing; J.R.D: data curation, investigation, methodology, validation, writing-review & editing; B.A.F.: investigation, methodology, validation, writing-review & editing; M.J.: data curation, investigation, methodology, validation, writing-review & editing; S.L.A.: investigation, funding acquisition, methodology, supervision, validation, writing-review & editing.

All authors gave final approval for publication and agreed to be held accountable for the work performed therein.

## Acknowledgements.

Thank you to an anonymous reviewer of Flanagan et al. 2021 for the original project idea. We also thank Estevan Gonzalez for assistance with copepod breeding and Dana Austria for preliminary respirometry work.

## Electronic Supplementary Material

**Figure S1.**
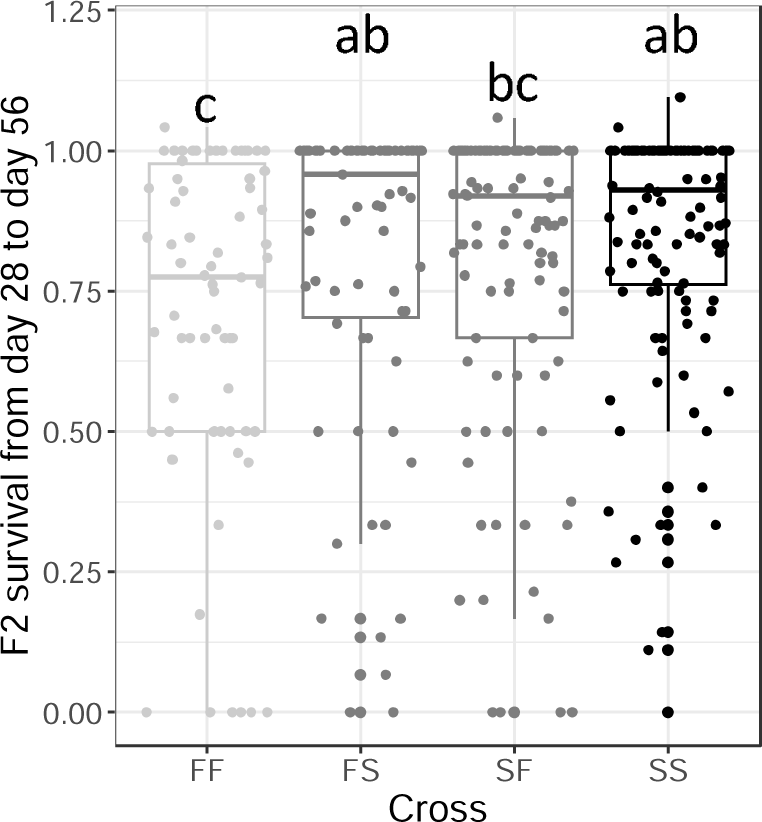
Proportion of F2 individuals surviving from day 28 to day 56. Letters above bars denote differences between cross types from pairwise Wilcoxon tests. *N* = 124-182 observations from 25-41 families for each cross.

**Figure S2.**
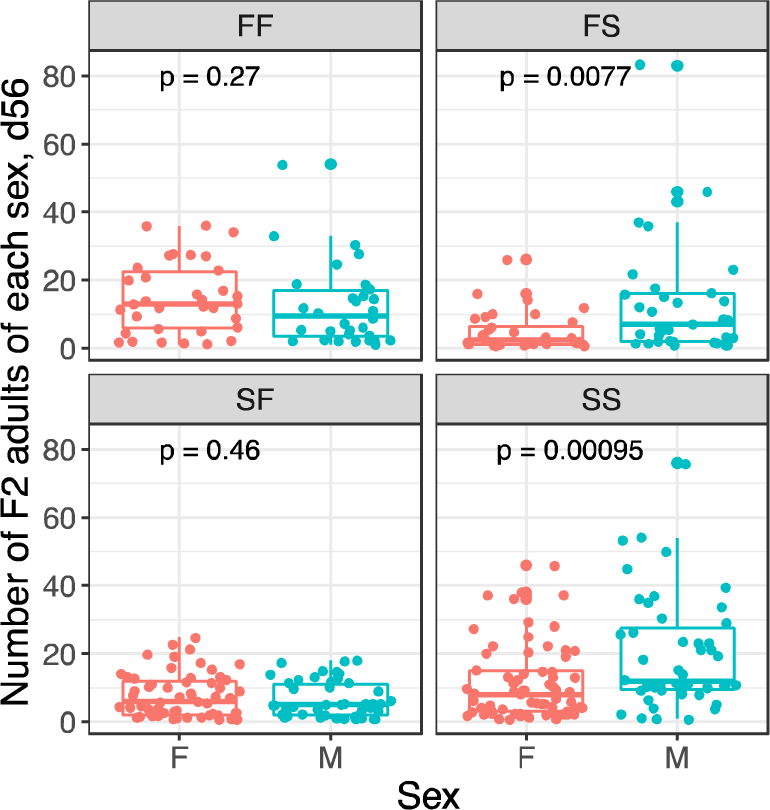
F2 sex ratio, measured as the number of females (F) and males (M) surviving to day 56 (per family per week). *P*-values indicate sex differences by pairwise Wilcoxon tests. *N* = 62-91 observations from 25-41 families for each cross.

**Figure S3.**
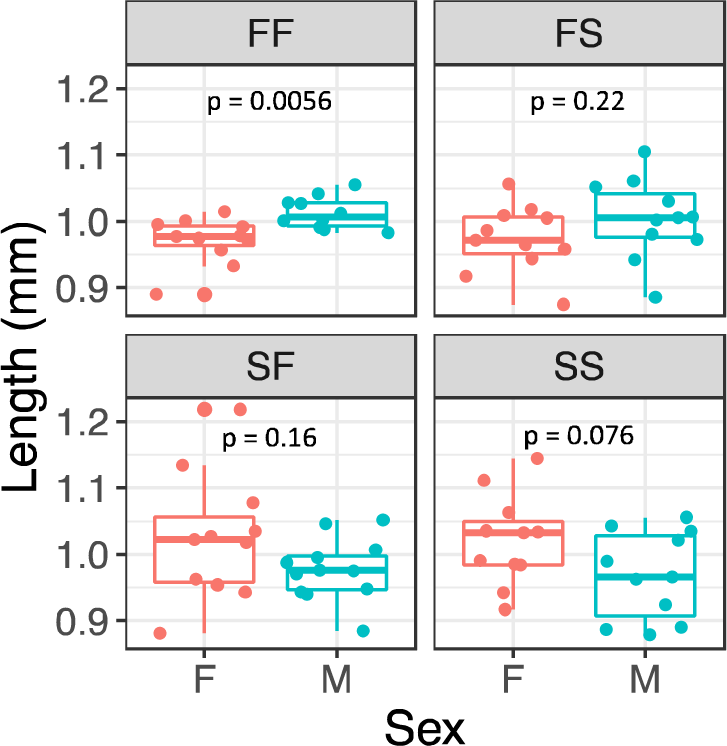
Body length in females (F) and males (M) in each F2 cross. *P*-values indicate sex differences by pairwise *t*-tests of log-transformed data. *N* 10-12 individuals per sex per cross.

**Figure S4.**
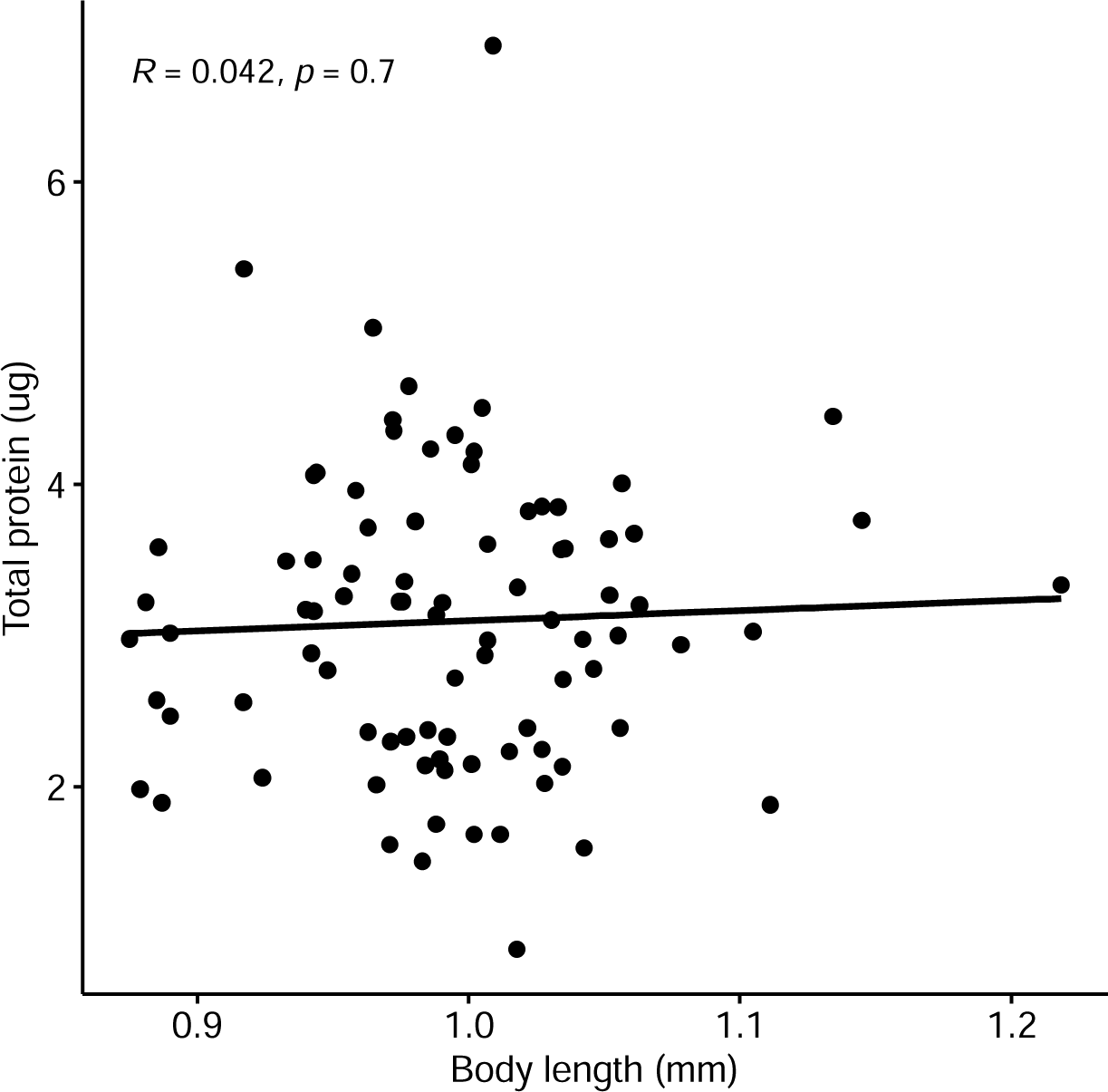
Correlation (Pearson’s) between body length and total protein content in F2 animals. *N* = 88 individuals.

## References

1. Boore, J. L. Animal mitochondrial genomes. Nucleic Acids Res. 27, 1767–1780 (1999).

2. Bar-Yaacov, D., Blumberg, A. & Mishmar, D. Mitochondrial-nuclear co-evolution and its effects on OXPHOS activity and regulation. Biochim. Biophys. Acta BBA - Gene Regul. Mech. 1819, 1107–1111 (2012).

3. Hill, G. E. Sex linkage of nuclear-encoded mitochondrial genes. Heredity 112, 469–470 (2014).

4. Burton, R. S. & Barreto, F. S. A disproportionate role for mtDNA in Dobzhansky-Muller incompatibilities? Mol. Ecol. 21, 4942–4957 (2012).

5. Aw, W. C., Garvin, M. R., Melvin, R. G. & Ballard, J. W. O. Sex-specific influences of mtDNA mitotype and diet on mitochondrial functions and physiological traits in Drosophila melanogaster. PLOS ONE 12, e0187554 (2017).

6. Wu, B. N. & O’Sullivan, A. J. Sex Differences in Energy Metabolism Need to Be Considered with Lifestyle Modifications in Humans. J. Nutr. Metab. 2011, 1–6 (2011).

7. Kyrgiafini, M.-A., Giannoulis, T., Moutou, K. A. & Mamuris, Z. Investigating the Impact of a Curse: Diseases, Population Isolation, Evolution and the Mother’s Curse. Genes 13, 2151 (2022).

8. Lichtenbelt, W. D. V. M., Wesselingh, R. A., Vogel, J. T. & Albers, K. B. M. Energy Budgets in Free-Living Green Iguanas in a Seasonal Environment. Ecology 74, 1157–1172 (1993).

9. Hutchings, J. A. Survival consequences of sex-biased growth and the absence of a growth-mortality trade-off. Funct. Ecol. 20, 347–353 (2006).

10. King, A. M., Kirkwood, T. B. L. & Shanley, D. P. Explaining sex differences in lifespan in terms of optimal energy allocation in the baboon: SEX ROLE SPECIALIZATION AND LIFESPAN. Evolution 71, 2280–2297 (2017).

11. Frank, S. & Hurst, L. Mitochondria and male disease. Nature 383, 224 (1996).

12. Innocenti, P., Morrow, E. H. & Dowling, D. K. Experimental Evidence Supports a Sex-Specific Selective Sieve in Mitochondrial Genome Evolution. Science 332, 845–848 (2011).

13. Nagarajan-Radha, V., Aitkenhead, I., Clancy, D. J., Chown, S. L. & Dowling, D. K. Sex-specific effects of mitochondrial haplotype on metabolic rate in Drosophila melanogaster support predictions of the Mother’s Curse hypothesis. Philos. Trans. R. Soc. B Biol. Sci. 375, 20190178 (2020).

14. Camus, M. F., Wolf, J. B. W., Morrow, E. H. & Dowling, D. K. Single Nucleotides in the mtDNA Sequence Modify Mitochondrial Molecular Function and Are Associated with Sex-Specific Effects on Fertility and Aging. Curr. Biol. 25, 2717–2722 (2015).

15. Carnegie, L., Reuter, M., Fowler, K., Lane, N., & M. Florencia Camus. Mother’s curse is pervasive across a large mito-nuclear Drosophila panel. http://biorxiv.org/lookup/doi/10.1101/2020.09.23.308791 (2020) doi:10.1101/2020.09.23.308791.

16. Mossman, J. A., Tross, J. G., Li, N., Wu, Z. & Rand, D. M. Mitochondrial-Nuclear Interactions Mediate Sex-Specific Transcriptional Profiles in Drosophila. Genetics 204, 613–630 (2016).

17. Mossman, J. A., Biancani, L. M., Zhu, C.-T. & Rand, D. M. Mitonuclear Epistasis for Development Time and Its Modification by Diet in Drosophila. Genetics 203, 463–484 (2016).

18. Watson, E. T., Flanagan, B. A., Pascar, J. A. & Edmands, S. Mitochondrial effects on fertility and longevity in Tigriopus californicus contradict predictions of the mother’s curse hypothesis. Proc. R. Soc. B Biol. Sci. 289, 20221211 (2022).

19. Beery, A. K. & Zucker, I. Sex bias in neuroscience and biomedical research. Neurosci. Biobehav. Rev. 35, 565–572 (2011).

20. Woitowich, N. C., Beery, A. & Woodruff, T. A 10-year follow-up study of sex inclusion in the biological sciences. eLife 9, e56344 (2020).

21. Edmands, S. Sex Ratios in a Warming World: Thermal Effects on Sex-Biased Survival, Sex Determination, and Sex Reversal. J. Hered. 112, 155–164 (2021).

22. Clancy, D. J. Variation in mitochondrial genotype has substantial lifespan effects which may be modulated by nuclear background. Aging Cell 7, 795–804 (2008).

23. Gonzalez-Freire, M. et al. Reconsidering the Role of Mitochondria in Aging. J. Gerontol. A. Biol. Sci. Med. Sci. 70, 1334–1342 (2015).

24. Clutton-Brock, T. H. & Isvaran, K. Sex differences in ageing in natural populations of vertebrates. Proc. R. Soc. B Biol. Sci. 274, 3097–3104 (2007).

25. Tower, J. Mitochondrial maintenance failure in aging and role of sexual dimorphism. Arch. Biochem. Biophys. 576, 17–31 (2015).

26. Tower, J. Sex-Specific Gene Expression and Life Span Regulation. Trends Endocrinol. Metab. 28, 735–747 (2017).

27. Committee on the Ethical and Social Policy Considerations of Novel Techniques for Prevention of Maternal Transmission of Mitochondrial DNA Diseases, Board on Health Sciences Policy, Institute of Medicine, & National Academies of Sciences, Engineering, and Medicine. Mitochondrial Replacement Techniques: Ethical, Social, and Policy Considerations. 21871 (National Academies Press, 2016). doi:10.17226/21871.

28. Claiborne, A. B., English, R. A. & Kahn, J. P. Finding an ethical path forward for mitochondrial replacement. Science 351, 668–670 (2016).

29. Babcock, C. S. & Asmussen, M. A. Effects of Differential Selection in the Sexes on Cytonuclear Polymorphism and Disequilibria. Genetics 144, 839–853 (1996).

30. Babcock, C. S. & Asmussen, M. A. Effects of Differential Selection in the Sexes on Cytonuclear Dynamics: Life Stages With Sex Differences. Genetics 149, 2063–2077 (1998).

31. Rand, D. M., Clark, A. G. & Kann, L. M. Sexually Antagonistic Cytonuclear Fitness Interactions in Drosophila melanogaster. Genetics 159, 173–187 (2001).

32. Jelić, M. et al. Sex-specific effects of sympatric mitonuclear variation on fitness in Drosophila subobscura. BMC Evol. Biol. 15, 135 (2015).

33. Rawson, P. D. & Burton, R. S. Functional coadaptation between cytochrome c and cytochrome c oxidase within allopatric populations of a marine copepod. Proc. Natl. Acad. Sci. 99, 12955–12958 (2002).

34. Willett, C. S. & Burton, R. S. Evolution of Interacting Proteins in the Mitochondrial Electron Transport System in a Marine Copepod. Mol. Biol. Evol. 21, 443–453 (2004).

35. Ellison, C. K. & Burton, R. S. Genotype-dependent variation of mitochondrial transcriptional profiles in interpopulation hybrids. Proc. Natl. Acad. Sci. 105, 15831–15836 (2008).

36. Foley, B. R., Rose, C. G., Rundle, D. E., Leong, W. & Edmands, S. Postzygotic isolation involves strong mitochondrial and sex-specific effects in Tigriopus californicus, a species lacking heteromorphic sex chromosomes. Heredity 111, 391–401 (2013).

37. Barreto, F. S. et al. Genomic signatures of mitonuclear coevolution across populations of Tigriopus californicus. Nat. Ecol. Evol. 2, 1250–1257 (2018).

38. Lima, T. G., Burton, R. S. & Willett, C. S. Genomic scans reveal multiple mito-nuclear incompatibilities in population crosses of the copepod Tigriopus californicus. Evolution 73, 609–620 (2019).

39. Healy, T. M. & Burton, R. S. Strong selective effects of mitochondrial DNA on the nuclear genome. Proc. Natl. Acad. Sci. 117, 6616–6621 (2020).

40. Burton, R. S. INTRASPECIFIC PHYLOGEOGRAPHY ACROSS THE POINT CONCEPTION BIOGEOGRAPHIC BOUNDARY. Evolution 52, 734–745 (1998).

41. Edmands, S. Heterosis and outbreeding depression in interpopulation crosses spanning a wide range of divergence. Evolution 53, 1757–1765 (1999).

42. Ar-Rushdi, A.H. The polygenic basis of sex-ratio in Tigriopus. Proc Int Congr Genet 10th 2, 8– 9.

43. Voordouw, M. & Anholt, B. Heritability of sex tendency in a harpacticoid copepod, Tigriopus californicus. Evolution 56, 1754–1763 (2002).

44. Alexander, H. J., Richardson, J. M. L. & Anholt, B. R. Multigenerational response to artificial selection for biased clutch sex ratios in Tigriopus californicus populations. J. Evol. Biol. 27, 1921–1929 (2014).

45. Alexander, H. J., Richardson, J. M. L., Edmands, S. & Anholt, B. R. Sex without sex chromosomes: genetic architecture of multiple loci independently segregating to determine sex ratios in the copepod Tigriopus californicus. J. Evol. Biol. 28, 2196–2207 (2015).

46. Willett, C. S. POTENTIAL FITNESS TRADE-OFFS FOR THERMAL TOLERANCE IN THE INTERTIDAL COPEPOD TIGRIOPUS CALIFORNICUS: TEMPERATURE ADAPTATION IN TIGRIOPUS CALIFORNICUS. Evolution 64, 2521–2534 (2010).

47. Kelly, M. W., Sanford, E. & Grosberg, R. K. Limited potential for adaptation to climate change in a broadly distributed marine crustacean. Proc. R. Soc. B Biol. Sci. 279, 349–356 (2012).

48. Foley, H. B. et al. Sex-specific stress tolerance, proteolysis, and lifespan in the invertebrate Tigriopus californicus. Exp. Gerontol. 119, 146–156 (2019).

49. Flanagan, B. A., Huang, E. & Edmands, S. Exogenous oxidative stressors elicit differing age and sex effects in Tigriopus californicus. Exp. Gerontol. 166, 111871 (2022).

50. Deconinck, A. & Willett, C. S. Hypoxia tolerance, but not low pH tolerance, is associated with a latitudinal cline across populations of Tigriopus californicus. PLOS ONE 17, e0276635 (2022).

51. Li, N., Arief, N. & Edmands, S. Effects of oxidative stress on sex-specific gene expression in the copepod Tigriopus californicus revealed by single individual RNA-seq. Comp. Biochem. Physiol. Part D Genomics Proteomics 31, 100608 (2019).

52. Li, N., Flanagan, B. A., Partridge, M., Huang, E. J. & Edmands, S. Sex differences in early transcriptomic responses to oxidative stress in the copepod Tigriopus californicus. BMC Genomics 21, 759 (2020).

53. Flanagan, B. A., Li, N. & Edmands, S. Mitonuclear interactions alter sex-specific longevity in a species without sex chromosomes. Proc. R. Soc. B Biol. Sci. 288, 20211813 (2021).

54. Reik, W. & Walter, J. Genomic imprinting: parental influence on the genome. Nat. Rev. Genet. 2, 21–32 (2001).

55. Ar-Rushdi, A.H. The cytology of achiasmatic meiosis in the female Tigriopus (Copepoda). Chromosoma 13, 1754–1763 (1963).

56. Burton, R.S., Feldman, M.W., & Swisher, S.G. Linkage relationships among five enzyme-encoding gene loci in the copepod Tigriopus californicus: a genetic confirmation of achiasmatic meiosis. Biochem Gen 19, 1237–1245 (1981).

57. R Core Team. R: A language and environment for statistical computing. (2022).

58. Rooney, J. P. et al. PCR Based Determination of Mitochondrial DNA Copy Number in Multiple Species. in Mitochondrial Regulation (eds. Palmeira, C. M. & Rolo, A. P.) vol. 1241 23–38 (Springer New York, 2015).

59. Schneider, C. A., Rasband, W. S. & Eliceiri, K. W. NIH Image to ImageJ: 25 years of image analysis. Nat. Methods 9, 671–675 (2012).

60. Schmidt-Nielsen, K. Scaling: Why is Animal Size So Important? (Cambridge University Press, 1984).

61. Prado-Cabrero, A., Herena-Garcia, R. & Nolan, J. M. Intensive production of the harpacticoid copepod Tigriopus californicus in a zero-effluent ‘green water’ bioreactor. Sci. Rep. 12, 466 (2022).

62. Burton, R. S. HYBRID BREAKDOWN IN DEVELOPMENTAL TIME IN THE COPEPOD TIGRIOPUS CALIFORNICUS. Evolution 44, 1814–1822 (1990).

63. Courret, C., Chang, C.-H., Wei, K. H.-C., Montchamp-Moreau, C. & Larracuente, A. M. Meiotic drive mechanisms: lessons from Drosophila. Proc. R. Soc. B Biol. Sci. 286, 20191430 (2019).

64. Ballard, J. W. O., Melvin, R. G., Miller, J. T. & Katewa, S. D. Sex differences in survival and mitochondrial bioenergetics during aging in Drosophila. Aging Cell 6, 699–708 (2007).

65. Yin, P. H. et al. Alteration of the copy number and deletion of mitochondrial DNA in human hepatocellular carcinoma. Br. J. Cancer 90, 2390–2396 (2004).

66. Mengel-From, J. et al. Mitochondrial DNA copy number in peripheral blood cells declines with age and is associated with general health among elderly. Hum. Genet. 133, 1149–1159 (2014).

67. Ashar, F. N. et al. Association of mitochondrial DNA levels with frailty and all-cause mortality. J. Mol. Med. 93, 177–186 (2015).

68. Kristensen, T. N., Loeschcke, V., Tan, Q., Pertoldi, C. & Mengel-From, J. Sex and age specific reduction in stress resistance and mitochondrial DNA copy number in Drosophila melanogaster. Sci. Rep. 9, 12305 (2019).

69. Dybdahl, M. F. Selection on life-history traits across a wave exposure gradient in the tidepool copepod Tigriopus californicus (Baker). J. Exp. Mar. Biol. Ecol. 192, 195–210 (1995).

70. Norin, T. & Metcalfe, N. B. Ecological and evolutionary consequences of metabolic rate plasticity in response to environmental change. Philos. Trans. R. Soc. B Biol. Sci. 374, 20180180 (2019).

71. Videlier, M., Careau, V., Wilson, A. J. & Rundle, H. D. Quantifying selection on standard metabolic rate and body mass in Drosophila melanogaster. Evolution 75, 130–140 (2021).

72. Ellison, C. & Burton, R. Disruption of mitochondrial function in interpopulation hybrids of Tigriopus californicus. Evolution 60, 1382–1391 (2006).

73. Olson, J. R., Cooper, S. J., Swanson, D. L., Braun, M. J. & Williams, J. B. The Relationship of Metabolic Performance and Distribution in Black-Capped and Carolina Chickadees. Physiol. Biochem. Zool. 83, 263–275 (2010).

74. Gvoždík, L. Metabolic costs of hybridization in Newts. Folia Zool. 61, 197–201 (2012).

75. Hoekstra, L. A., Siddiq, M. A. & Montooth, K. L. Pleiotropic Effects of a Mitochondrial– Nuclear Incompatibility Depend upon the Accelerating Effect of Temperature in Drosophila. Genetics 195, 1129–1139 (2013).

76. McFarlane, S. E., Sirkiä, P. M., Ålund, M. & Qvarnström, A. Hybrid Dysfunction Expressed as Elevated Metabolic Rate in Male Ficedula Flycatchers. PLOS ONE 11, e0161547 (2016).

77. Davies, R. et al. Hybridization in Sunfish Influences the Muscle Metabolic Phenotype. Physiol. Biochem. Zool. 85, 321–331 (2012).

78. Barreto, F. S. & Burton, R. S. Elevated oxidative damage is correlated with reduced fitness in interpopulation hybrids of a marine copepod. Proc. R. Soc. B Biol. Sci. 280, 20131521 (2013).

79. Moreno-Loshuertos, R. et al. Differences in reactive oxygen species production explain the phenotypes associated with common mouse mitochondrial DNA variants. Nat. Genet. 38, 1261–1268 (2006).

80. Gusdon, A. M., Votyakova, T. V., Reynolds, I. J. & Mathews, C. E. Nuclear and Mitochondrial Interaction Involving mt-Nd2 Leads to Increased Mitochondrial Reactive Oxygen Species Production. J. Biol. Chem. 282, 5171–5179 (2007).

81. Lane, N. Mitonuclear match: Optimizing fitness and fertility over generations drives ageing within generations: Problems and Paradigms. BioEssays 33, 860–869 (2011).

82. Đorđević, M., et al. Sex-specific mitonuclear epistasis and the evolution of mitochondrial bioenergetics, ageing, and life history in seed beetles: SEX-SPECIFIC MITONUCLEAR EPISTASIS. Evolution 71, 274–288 (2017).

